# COP1 promotes seed germination by destabilizing RGA-LIKE2 (RGL2) in Arabidopsis

**DOI:** 10.1101/2021.12.01.470837

**Authors:** Byoung-Doo Lee, Yehyun Yim, Esther Cañibano, Suk-Hwan Kim, Marta García-León, Vicente Rubio, Sandra Fonseca, Nam-Chon Paek

## Abstract

Under favorable moisture, temperature and light conditions, gibberellin (GA) biosynthesis is induced and triggers seed germination. A major mechanism by which GA promotes seed germination is by promoting the degradation of the DELLA protein RGL2, a major repressor of germination in Arabidopsis seeds. Analysis of seed germination phenotypes of *constitutively photomorphogenic 1* (*cop1*) mutants and complemented *COP1-OX/cop1-4* lines in response to GA and paclobutrazol (PAC) suggested a positive role for COP1 in seed germination and a relation with GA signaling. *cop1-4* mutant seeds showed PAC hypersensitivity, but transformation with a *COP1* overexpression construct rendered them PAC insensitive, with a phenotype similar to that of *rgl2* mutant (*rgl2-SK54*) seeds. Furthermore, *cop1-4 rgl2-SK54* double mutants showed a PAC-insensitive germination phenotype like that of *rgl2-SK54*, identifying COP1 as an upstream negative regulator of RGL2. COP1 interacts directly with RGL2 and *in vivo* this interaction is strongly enhanced by SPA1. COP1 directly ubiquitinates RGL2 to promote its degradation. Moreover, GA stabilizes COP1 with consequent RGL2 destabilization. By uncovering this COP1-RGL2 regulatory module, we reveal a novel mechanism whereby COP1 positively regulates seed germination and controls the expression of germination-promoting genes.

## Introduction

In its seed form, a new plant generation can be dispersed and then wait to develop and grow once favorable environmental conditions, such as optimal moisture, light, and temperature, exist (Bewley, 1997). Favorable conditions trigger changes in the phytohormone levels of mature seeds, including a decrease in abscisic acid (ABA) and an increase in gibberellins (GA), which together, inhibit dormancy and promote seed germination (Steber *et al*., 1998; Holdsworth *et al*., 2008). Seed imbibition induces GA biosynthesis and triggers germination. Arabidopsis (*Arabidopsis thaliana*) mutants with impaired GA biosynthesis (e.g., *ga1-3* and *ga2*) fail to germinate in the absence of exogenous GA, underlining its importance for germination (Koornneef & van der Veen, 1980).

GA is perceived by the receptor protein GA-INSENSITIVE DWARF 1 (GID1), which then changes conformation and binds to DELLA proteins, central repressors in the GA signaling pathway (Ueguchi-Tanaka *et al*., 2005; Murase *et al*., 2008; Shimada *et al*., 2008). The formation of the GID1-GA-DELLA complex triggers GA-mediated DELLA degradation by the F-box protein SLEEPY1 (SLY1; SCF^SLY1^ complex) and its homolog SNEEZY1/SLY2 through ubiquitin-dependent proteolysis, and induces the expression of GA-responsive genes (McGinnis *et al*., 2003; Dill *et al*., 2004; Fu *et al*., 2004; Ariizumi *et al*., 2011). Thus, GA lifts DELLA repression of its downstream targets, triggering GA-mediated responses (Sun *et al*., 2010).

In Arabidopsis, five DELLA repressors, such as GA INSENSITIVE (GAI), REPRESSOR OF *ga1-3* (RGA), RGA-LIKE 1 (RGL1), RGL2, and RGL3, play overlapping yet distinct roles in many developmental processes, such as seed germination, stem elongation, and transition to flowering (Peng *et al*., 1997; Dill & Sun 2001; Lee *et al*., 2002; Wen & Chang, 2002; Cao *et al*., 2005). GAI and RGA are both involved in seed germination (Oh *et al*., 2007), but RGL2 is the major regulator of GA-mediated seed germination among the DELLA proteins. Under GA-deficient conditions, such as in the *ga1-3* mutant background or under paclobutrazol (PAC) treatment, only *rgl2* mutation, but not *rgl1* or *rgl3*, rescued seed germination rates under light conditions (Lee *et al*., 2002; Tyler *et al*., 2004; Cao *et al*., 2005).

The repressive role of RGL2 in seed germination operates through different molecular mechanisms that integrate GA, ABA, and light signals. RGL2 associates with transcription factors to regulate gene expression and control germination (Piskurewicz *et al*., 2008; Liu *et al*., 2016; Ravindran *et al*., 2017; Yang *et al*., 2019; Sanchez-Montesino *et al*., 2019). Furthermore, RGL2 represses the expression of the *GA-STIMULATED ARABIDOPSIS6* (*GASA6*), which is an integrator of glucose, GA and ABA signals in seed germination (Zhong *et al*., 2015). GASA6 locates mainly in cell wall and regulates cell wall loosening to favor cell elongation on embryonic axis during seed germination through EXPANSIN 1 (EXPA1) (Zhong *et al*., 2015).

Light impact on seed germination relies in part on phytochrome B (phyB) activation/inactivation by red/far-red light. Once activated, phyB promotes the degradation of PHYTOCHROME-INTERACTING FACTOR 1 (PIF1), a strong repressor of light-mediated seed germination (Oh *et al.,* 2004; Oh *et al*., 2006). Thus, *pif1* mutants display high germination rates in the dark (Oh et al., 2004). PIF1 regulate germination in many ways, by directly binds to the promoters of the DELLA repressors *GAI* and *RGA* to activate their expression and repress GA signaling (Oh *et al*., 2007); by directly repressing genes involved in GA biosynthesis and activating GA catabolism (Kim *et al*., 2016; Oh *et al*., 2007; Oh *et al*., 2009); by acting cooperatively with ABI3 in the dark to activate *SOMNUS* (*SOM*), a key negative regulator of seed germination; and by directly binding to the *ABI5* promoter (Park *et al.,* 2011; Kim *et al*., 2008). Thus, stabilization of PIF1 in the dark is a key to halting seed germination, as it mediates the repression and activation of the GA and ABA cascades, respectively (Oh *et al*., 2006, 2007; Kim *et al*., 2008).

Previous reports have shown that additional regulators of light signaling are involved in the control of seed germination. Among them is CONSTITUTIVELY PHOTOMORPHOGENIC 1 (COP1), a master regulator of photomorphogenesis. COP1, together with the E2 ubiquitin variant COP10, DE-ETIOLATED 1 (DET1), and COP9 SIGNALOSOME (CSN) proteins, is a member of the COP/DET/FUSCA family, whose strong mutants produce dark-purple-pigmented seeds and exhibit seedling-lethal phenotypes (McNellis *et al*., 1994). COP1, in association with SPA proteins, acts as part of a substrate adaptor module within CULLIN4 (CUL4)-based E3 ubiquitin ligases. The formation of CUL4^COP1-SPA^ complexes mediates the targeted degradation of many positive regulators of photomorphogenesis in darkness, as well as of other regulators of circadian clock and photoperiodic flowering (Lau & Deng, 2012). In the case of seed germination, the CUL4^COP1– SPA^ E3 ubiquitin ligase is necessary for the light-induced degradation of PIF1 in Arabidopsis. Accordingly, *cop1* and *spaQ* mutants display reduced seed germination in response to light, consistent with the higher abundance of PIF1 in these mutants compared to the wild-type (Zhu *et al*., 2015). Strikingly, in the dark, PIF1 acts as a cofactor of CUL4^COP1–SPA^ E3 ubiquitin ligase, enhancing its function to synergistically degrade HY5 and repress photomorphogenesis, being then degraded in the light (Xu *et al*., 2014; Zhu *et al*., 2015). Despite its role as a positive regulator of photomorphogenesis, HY5 binds to the *ABI5* promoter and is required for *ABI5* expression in developing seeds, positively controlling seed maturation and dormancy (Chen *et al*., 2008). Both COP1 and HY5 proteins have been recently shown to be involved in ABA-mediated inhibition of post-germination seedling development (Yadukrish *et al*., 2020a; Yadukrish *et al*., 2020b).

Here, we provide evidence that COP1 acts as a positive regulator of seed germination by limiting the accumulation of the DELLA protein RGL2. In germination analysis, the *cop1-4* weak mutant showed a PAC-hypersensitive germination phenotype, but the *COP1*-overexpressing plants exhibited PAC insensitivity in an RGL2-dependent manner. COP1 was stabilized by GA and directly interacted with and ubiquitinated RGL2 to promote its degradation, releasing the expression of downstream regulators of seed germination (such as *GASA6* and *EXPA1*). Taken together, our data suggest that COP1- and GA-mediated seed germination converges on RGL2 regulation. This finding contributes to the understanding of the regulatory role of COP1 in the germination of Arabidopsis seeds.

## Results

### COP1 is a regulator of seed germination

*COP1* is a member of the *COP/DET/FUS* gene family, whose strong mutants produce dark-purple-pigmented (fusca) seeds and exhibit a seedling-lethal phenotype (McNellis *et al*., 1994). We found that strong mutants of *COP1*, especially the *cop1-5* seedling-lethal mutant caused by a T-DNA insertion (Fig. **1A**), exhibited extremely delayed or failed seed germination (McNellis *et al*., 1994). To determine whether fusca phenotypes were in general associated to seed germination defects we tested the germination of another fusca mutant, *cop10-1*. COP10 belongs to the CDDD complex that conforms an E3 ligase (Lau and Deng 2012). The purple-colored seeds of the *cop10-1* seedling-lethal mutant (Fig. **1B**) (Wei *et al*., 1994) germinated normally, like wild-type seeds, under normal conditions (mock), indicating that the fusca phenotypes are not intrinsically associated with germination defects (Fig. **1C**). In the presence of 10 μM gibberellic acid (GA_3_), the germination rate of *cop1-5* seeds was greatly enhanced compared to that of mock-treated seeds (Fig. **1D**).

**Figure 1.**
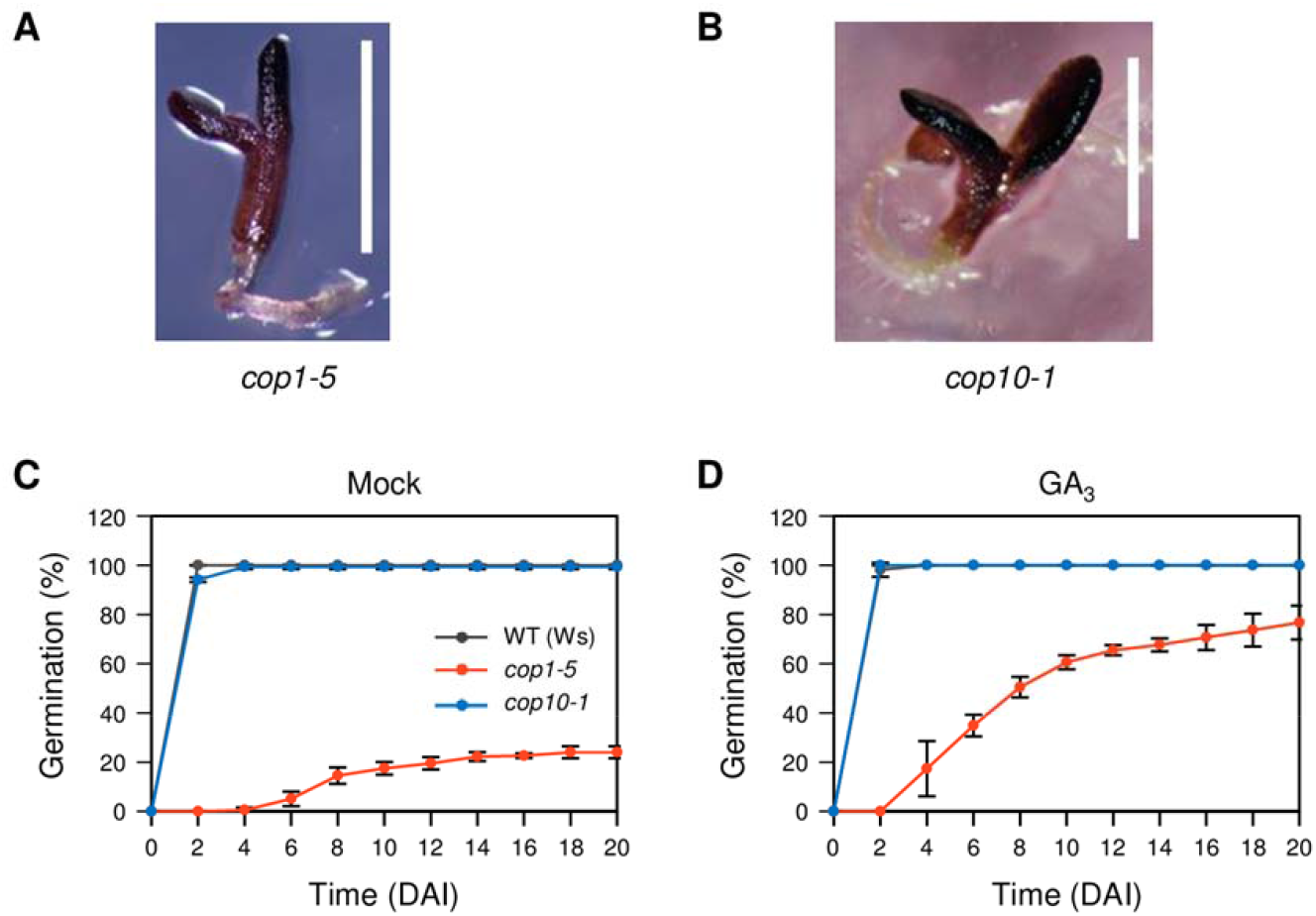
Germination rate of *cop1-5* seeds is dramatically enhanced in the presence of GA. (A, B) The seedling-lethal phenotypes of *cop1-5* (8 DAI) (A) and *cop10-1* (4 DAI) (B) mutants. (C, D) Germination rates of wild-type (WT; Ws ecotype), *cop1-5*, and *cop10-1* seeds on MS medium (mock) (C) or on the same medium containing 10 μM GA_3_ (D). Germination rates were determined by counting the seeds with protruding radicles over 20 days. The mean and SD were obtained from three biological repeats (∼100 seeds/repeat). The experiments were repeated five times with similar results. DAI, day(s) after incubation to 22°C under long days; GA, gibberellin.

To further confirm the regulatory role of COP1 in seed germination, we generated *COP1-OX/cop1-4* transgenic plants by transforming *cop1-4* (*cop1* weak allele) mutant plants (McNellis *et al*., 1994) with a *35S::COP1-GFP* construct (see Methods; Fig. **S1**). Next, we examined the germination rates of *cop1-4* and *COP1-OX/cop1-4* seeds in the presence of GA or the GA biosynthesis inhibitor paclobutrazol (PAC) (Fig. **2**). We found that the germination of *cop1-4* seeds was slightly, but significantly, delayed compared to that of wild-type seeds under normal conditions (mock treatment) but was fully restored in the presence of GA, (Fig. **2A,B**). Moreover, compared to wild-type seeds, the faster germination rate of *COP1-OX/cop1-4* seeds at 1.5 d after incubation (DAI) at 22°C in long day conditions (Fig. **2B**) indicated not only the full rescue of the *cop1-4* defect in delayed germination but also a positive effect of COP1 on seed germination. However, in the presence of PAC at 3 DAI (Fig. **2B,C**), the germination of *cop1-4* mutant seeds was impaired, whereas *COP1-OX/cop1-4* seeds germinated much faster and in higher percentage than wild-type seeds, showing strong insensitivity to the negative effect of PAC on seed germination. Thus, COP1’s effect on seed germination is stronger when GA levels are reduced. These results suggest that COP1 positively regulates seed germination, either by targeting a component of the GA signaling pathway involved in germination or by affecting a GA-independent pathway concurrently implicated in seed germination.

**Figure 2.**
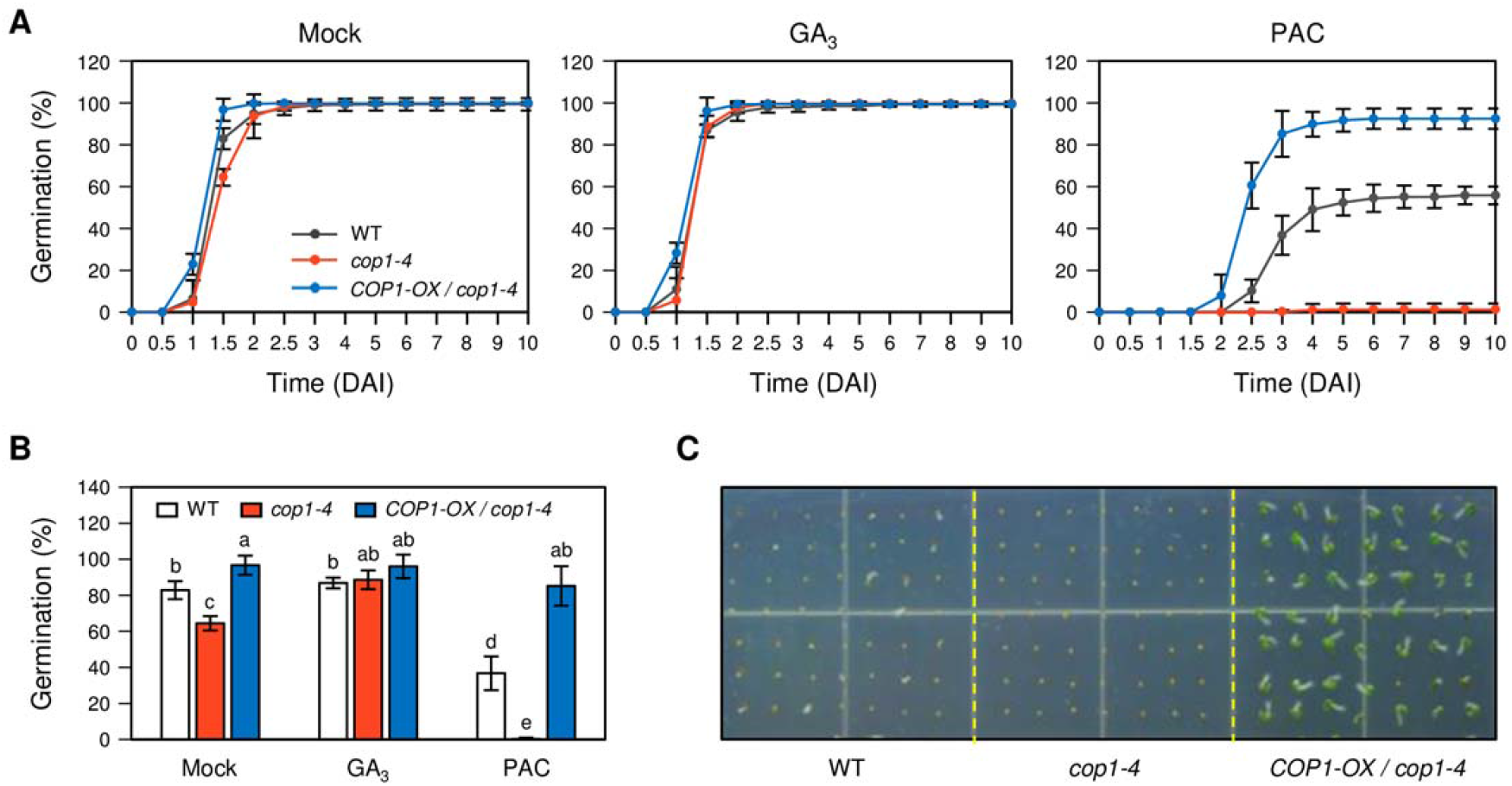
COP1 acts as a positive regulator in seed germination. (A) Germination rates of WT (Col-0 ecotype), *cop1-4*, and *COP1-OX/cop1-4* seeds on MS phytoagar medium (mock) or on the same medium containing 10 μM GA_3_ or 10 μM PAC from 0 to 10 DAI. (B) Germination scored at 1.5 DAI of the same lines and treatments as in (A). Germination rates were determined by counting the seeds with protruding radicles. The mean and SD were obtained from three independent repeats (∼100 seeds/repeat). Different letters indicate significantly different values according to a one-way ANOVA and Duncan’s least significant range test (*P* < 0.05). (C) Germination phenotypes of WT, *cop1-4*, and *COP1-OX/cop1-4* seeds on MS medium containing 10 μM PAC at 3 DAI. The experiments were repeated five times with similar results. DAI, day(s) after incubation to 22°C under LD; GA, gibberellin; PAC, paclobutrazol.

### COP1 upregulates the expression of germination-associated genes in imbibed seeds

During seed imbibition, GA upregulates the expression of downstream genes that induce cell wall remodeling needed for germination, expansins (EXPAs) and xyloglucan endotransglucosylases/endohydrolases (XTHs) (Weitbrecht *et al*., 2011; Shi *et al*., 2013). To examine the positive role of COP1 in seed germination at the transcriptional level, we analyzed the expression levels of four cell wall remodeling genes by qRT-PCR in *cop1-4* and *COP1-OX/cop1-4* seeds, as compared to wild-type seeds, at 3 DAI under normal conditions and in the presence of PAC (Fig. **3**). We found that expression of *EXPA1*, *EXPA2*, and *XTH33* was downregulated in the *cop1-4* seeds and only *XTH33* was upregulated in *COP1*-OX/*cop1-4* seeds under normal conditions. In the presence of PAC, the expression of *EXPA1, EXPA2, EXPA8* and *XTH33* was downregulated in *cop1-4* seeds and upregulated in *COP1-OX/cop1-4* seeds. Therefore, our results show that COP1 positively regulates the expression of genes involved in cell wall remodeling and that this regulatory role is evidenced upon inhibition of GA biosynthesis by PAC treatment, in accordance with the germination phenotypes observed in Fig**. 1**.

**Figure 3.**
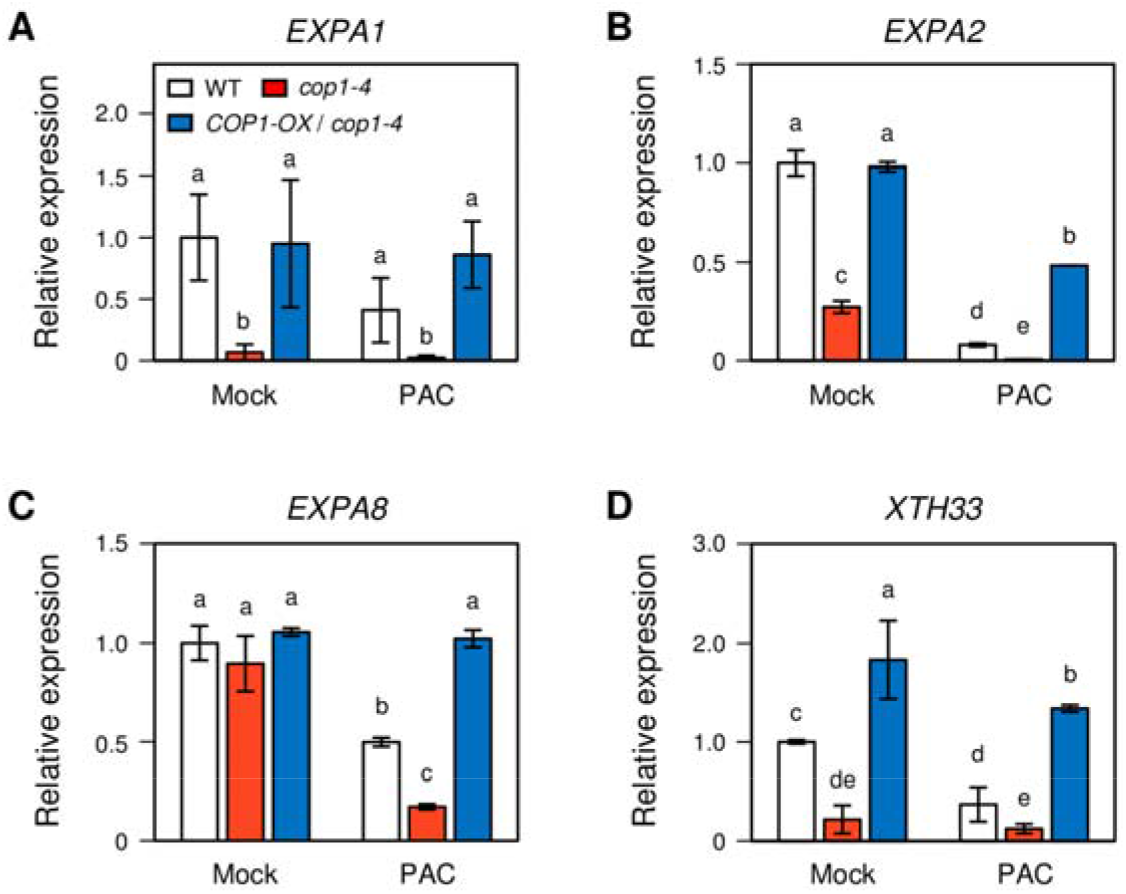
Expression profiles of germination-associated genes in *cop1-4* and *COP1-OX/cop1-4* seeds. (A-D) Expression levels of germination-associated genes, *EXPA1, EXPA2*, *EXPA8*, and *XTH33*, in *cop1-4* and *COP1-OX/cop1-4* seeds relative to those in WT seeds. After stratification, seeds were transferred to 22°C for germination and grown in MS phytoagar (mock) or in the presence of 10 μM PAC until 3 DAI, and analyzed by qRT-PCR. The expression level of each gene was normalized to that of *ACTIN2* (*ACT2*). Expression levels of each gene are shown relative to the expression of WT in the mock-treatment group, which is set as 1. The mean and SD were obtained from three biological repeats (∼30 seeds/repeat). Different letters indicate significantly different values according to a one-way ANOVA and Duncan’s least significant range test (*P* < 0.05). The experiments were repeated three times with similar results. DAI, day(s) after incubation to 22°C under LD; PAC, paclobutrazol.

### Light signaling regulators PIF1 and HY5 destabilized by COP1 do not explain the PAC hypersensitivity of *cop1* mutant seeds

Previous studies have shown that COP1 mediates PIF1 destabilization upon dark-to-light transition to allow the establishment of the photomorphogenic developmental program (Zhu *et al*. 2015). PIF1 also acts as a negative regulator of seed germination by promoting the expression of *DELLA*, *DOF AFFECTING GERMINATION 1* (*DAG1*), and *SOM* (Oh *et al*., 2007; Oh *et al*., 2009; Kim *et al*., 2008; Gabriele *et al*., 2010; Seo *et al*., 2009). However, a relation between COP1 and PIF1 in seed germination was never found. In addition, HY5, a positive regulator of photomorphogenesis that is destabilized by COP1 in darkness (Osterlund *et al*., 2000), is also involved in the ABA signaling pathway, where it negatively regulates seed germination (Chen *et al*., 2008; Yang *et al*., 2018). Because these studies do not discard a role of HY5 in GA-mediated seed germination, it could be speculated that HY5 destabilization by COP1 may be a mechanism to promote seed germination. To examine whether *HY5* and *PIF1* play a role in GA-mediated seed germination, we analyzed the germination rates of *pif1-1* and *hy5-215* single mutants and introgressed in a *cop1-4* background under normal conditions and in the presence of PAC (Fig. **S2 and S3**). Seeds of *pif1-1* and *pif1-1 cop1-4* mutants, when exposed to light, behaved similarly to wild-type and *cop1-4* seeds respectively, in mock and under PAC treatment, suggesting that *PIF1* is not involved in GA-mediated seed germination neither acts in the same pathway than *COP1* (Fig. **S2**). The analysis of *hy5-215* and *cop1-4 hy5-215* germination shows that *hy5* germinates slightly early in PAC regarding the WT and the same is true for *cop1-4 hy5-215* regarding *cop1-4* background, showing that in PAC *hy5* mutation contributes to a slight partial suppression of the *cop1-4* germination defects (Fig. **S3**). Altogether, these results suggested that neither PIF1 nor HY5 can be responsible for the strong *cop1-4* germination defects observed in the presence of PAC.

### RGL2, a negative regulator of GA-mediated seed germination, is epistatic to COP1

DELLA proteins are major negative regulators in the GA signaling pathway that comprise five proteins, encoded by *GAI*, *RGA*, *RGA-LIKE1* (*RGL1*), *RGL2*, and *RGL3* (Sun, 2010). All five DELLA proteins are stabilized by PAC as it represses GA biosynthesis. Among them, RGL2 is a key regulator of GA-mediated seed germination (Lee *et al*., 2002). When we examined germination rates of mutants of these five genes, only the *rgl2-SK54* mutant showed a PAC-insensitive phenotype, as previously reported (Fig. **S4**) (Lee *et al*., 2002; Tyler *et al*., 2004). To determine the genetic interaction between *COP1* and *RGL2* in seed germination, we generated *cop1-4 rgl2-SK54* double mutants. In addition, we obtained *COP1-OX/rgl2-SK54* transgenic plants by transforming *35S::COP1-GFP* into the *rgl2-SK54* mutant. Under normal conditions, *rgl2-SK54* single- and *cop1-4 rgl2-SK54* double-mutant seeds germinated faster than wild-type and *cop1-4* seeds (Fig. **4A****, left**). In the presence of PAC, however, *cop1-4 rgl2-SK54* seeds showed an almost PAC-insensitive phenotype, similar to that of *rgl2-SK54* seeds, indicating that the *rgl2* mutation suppresses the PAC-hypersensitive phenotype of *cop1-4* (Fig. **4A****, right**).

**Figure 4.**
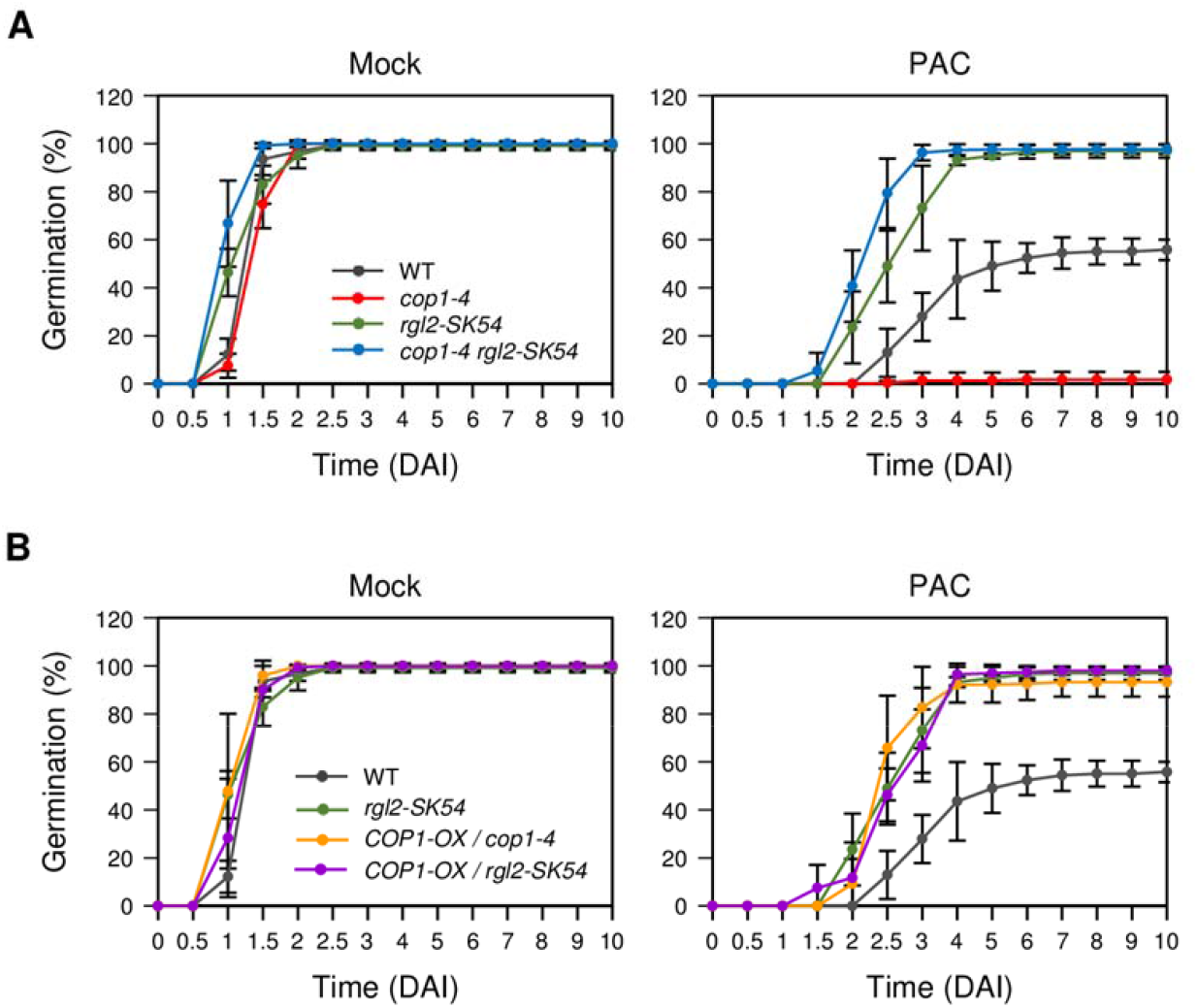
COP1 acts as an upstream regulator of RGL2 in seed germination. (A) Germination rates of WT (Col-0), *cop1-4, rgl2-SK54*, and *cop1-4 rgl2-SK54* seeds on MS phytoagar medium (mock). (B) Germination rates of WT (Col-0), *rgl2-SK54, COP1-OX/cop1-4*, and *COP1-OX/rgl2-SK54* seeds in MS phytoagar medium containing 10 μM PAC. After 3 days of stratification, the seeds were transferred to 22°C and the germination rate was determined by counting the seeds with protruding radicles at the indicated time from 0 to 10 DAI. The mean and SD were obtained from three biological repeats (∼100 seeds/repeat). The experiments were repeated three times with similar results. DAI, day(s) after incubation to 22°C under LD; PAC, paclobutrazol.

*COP1-OX/cop1-4* seeds germinated much faster and in higher percentage than wild-type seeds, and similarly to *rgl2-SK54* and *COP1-OX/rgl2-SK54* seeds, in the presence of PAC, suggesting that RGL2 acts downstream of COP1 and could be a COP1 target (Fig. **4B****, right**). Following up on the idea that an active COP1 should be then necessary to degrade RGL2 and promote seed germination, we measured the germination rates of the *cop1-4* lines overexpressing *COP1* variants that failed to dimerize or enter the nucleus, thus compromising COP1 E3 ligase function (Lee *et al*. 2017; Stacey *et al*., 1999). COP1^WT^-GFP and COP1^L105A^-GFP fusions, which are able to form COP1 homodimers, fully complemented the *cop1-4* germination defects in the presence of PAC, whereas COP1^L170A^-GFP, whose ability to form dimers is severely impaired, and COP1^cyt^-GFP, which is retained in the cytoplasm, did not rescue the *cop1-4* germination defects (Fig. **S5**). Although we could not rule out the possibility that these effects might be due to collateral effects of *cop1* mutation, these results are consistent with the fact that COP1 dimerization is required for target degradation, suggesting that the role of COP1 in seed germination requires a fully active nuclear E3 ubiquitin ligase activity.

### COP1 interacts with and destabilizes RGL2

We then investigated whether COP1 regulates RGL2 indirectly at the transcriptional level or directly at the post-translational level. For this, we analyzed the expression levels of *RGL2* and *COP1* by qRT-PCR in *cop1-4* and *rgl2-SK54* seeds at 3 DAI, respectively, and compared them to those in wild-type seeds. We found that the expression levels of *RGL2* in *cop1-4* and those of *COP1* in *rgl2-SK54* did not differ significantly from those of the wild-type (Fig. **S6A,B**). These results indicated that COP1 does not regulate *RGL2* at the transcriptional level.

Next, to examine whether COP1 interacts with and destabilizes RGL2 post-translationally, we first tested the physical interaction between the two proteins. Y2H assays revealed that, whereas full-length COP1 did not interact with RGL2, a truncated version of COP1 containing the RING domain could directly interact with RGL2 (Fig. **5A**). Since all COP1 fragments and RGL2 are being expressed in yeast (Fig. **S7**), it seems that the full-length COP1 is less efficient in promoting a direct interaction than the COP1 RING domain alone in the yeast cells. To overcome this technical limitation, we performed *in vitro* pull-down assays with recombinant proteins expressed in *E. coli* (Fig. **5B**). *In vitro* purified MBP-COP1 could pull-down GST-RGL2 fusions, thus supporting a direct interaction between the full-length COP1 and RGL2. To further confirm this interaction we performed semi-*in vivo* pull-down assays in the opposite direction. In this experiment, MBP-RGL2 could pull-down COP1-GFP from Arabidopsis seedling extracts whereas MBP protein alone could not (Fig. **5C**).

**Figure 5.**
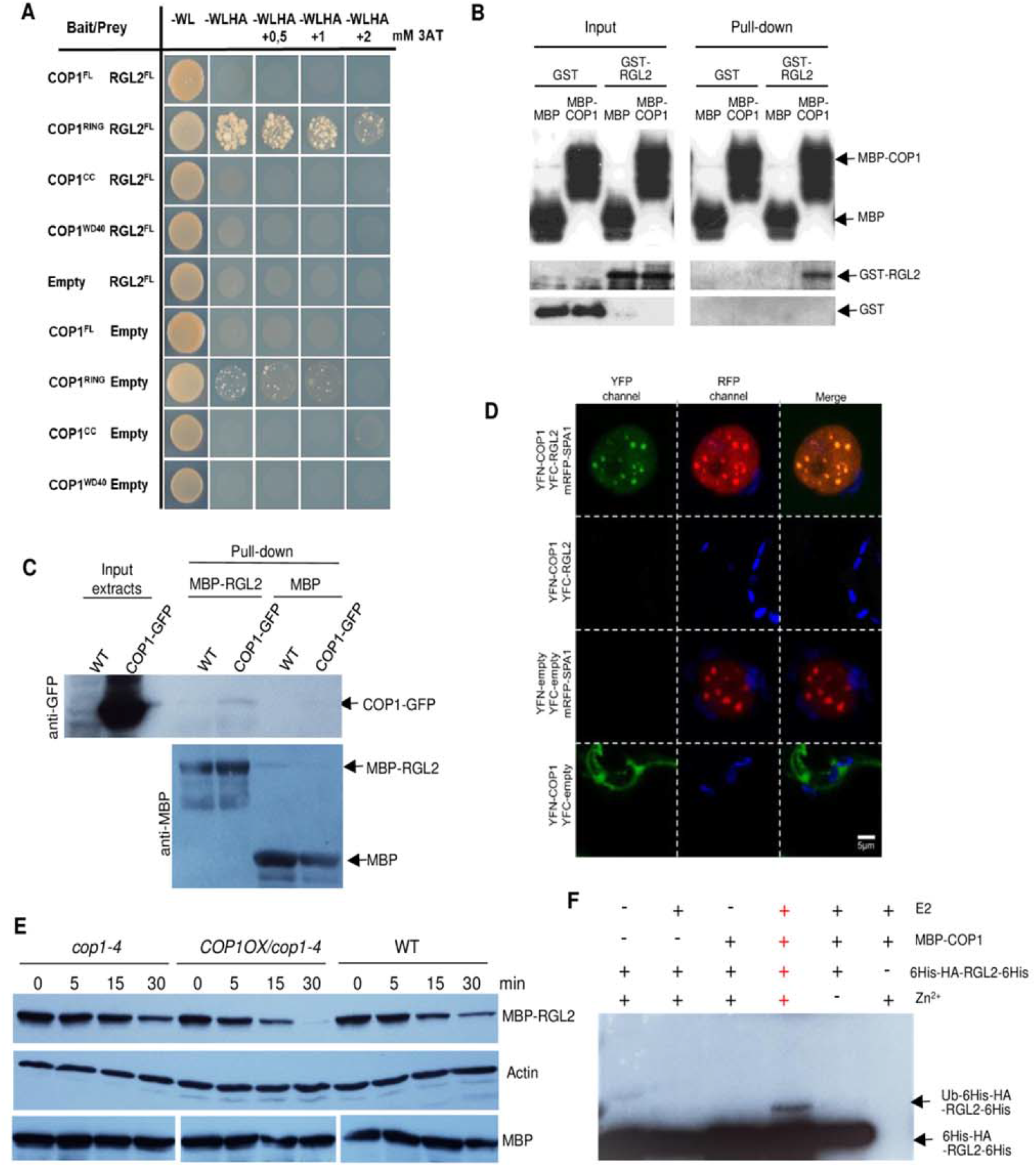
COP1 interacts with and destabilizes RGL2. (A) Interaction of COP1 and RGL2 in Y2H assays. Full-length (FL) COP1, as well as three of its individual domains (RING, coiled-coil (CC), and WD40-repeat (WD40)), were used as baits and full-length RGL2 as prey. Selection for interaction was performed in selective media containing increasing concentrations of 3-Amino-1,2,4-Triazol (3-AT). (B) MBP-COP1 pulls-down GST-RGL2 *in vitro*. Fusion proteins were detected with an anti-MBP antibody and anti-GST antibody. (C) Semi *in vivo* pull-down assays. GST-RGL2 pulls down COP1-GFP from Arabidopsis protein extracts. (D) BiFC assay showing that COP1 and RGL2 interact in the presence of SPA1. The indicated constructs were expressed in *N. benthamiana* leaves and observed by confocal microscopy. One representative nucleous is shown. (E) Degradation of MBP-RGL2 fusion by soluble protein extracts from seedlings of different genotypes as determined with a cell-free degradation assay. MBP-RGL2 proteins were incubated for the indicated times (min) with total soluble protein extracted from 6-d-old WT (Col-0), *cop1-4*, or *COP1-OX/cop1-4* etiolated seedlings. As a control for equivalent extract amounts actin is shown.MBP when expressed alone remained stable. An anti-MBP antibody was used for fusion protein-detection. Protein levels at each time point are shown relative to those in the input sample (time 0) and normalized to the corresponding actin loading, and set as 1. The experiments were repeated three times with similar results. (F) COP1 ubiquitinates RGL2 *in vitro*. RGL2 (6His-HA-RGL2-6His fusion) ubiquitination assays were performed using MBP-COP1 (or MBP MBP-COP1 without Zn^2+^ as a negative control), rice E2 Rad6 (E2), and yeast E1 (E1; Boston Biochem). Ubiquitinated RGL2 was detected using an anti-HA antibody.

In addition, to confirm whether this interaction occurs in vivo we performed bimolecular fluorescent complementation (BiFC) assays in *N. benthamiana* leaves. Contrary to the *in vitro* results, the co-expression of truncated YFN-COP1 and truncated YFC-RGL2 constructs showed no YFP (Yellow Fluorescent Protein) reconstitution signal. Based on the recent results by Blanco-Touriñan et al., (2020) where the addition of SPA1 was necessary to visualize the interaction between COP1 and the DELLA proteins RGA and GAI, we tested the effect of mRFP-SPA1 addition to COP1-RGL2 BiFC assays. Co-expression with SPA1 fusion rendered a very strong YFP-reconstitution signal visible in nuclear speckles, suggesting that SPA1 protein is necessary for the *in vivo* efficient recognition of RGL2 by COP1 (Fig. **5D**).

To examine whether COP1 destabilizes RGL2 *in vivo*, we examined the changes in RGL2 levels in the *cop1-4* and *COP1-OX/cop1-4* backgrounds using cell-free (or *in vitro*) degradation assays. To this end, we incubated MBP-RGL2 protein fusions with the total soluble protein extracts of wild-type, *cop1-4*, and *COP1-OX/cop1-4* grown in the dark and detected the changes in MBP-RGL2 levels over 30 min by immunoblot analysis using an anti-MBP antibody. MBP-RGL2 was degraded faster in *COP1-OX/cop1-4* extracts and slowly in *cop1-4* extracts than in wild-type whereas MBP alone remained stable (Fig. **5E**). Furthermore, *in vitro* ubiquitination assays showed that RGL2 (6His-HA-RGL2-6His fusion) was directly ubiquitinated by MBP-COP1 (Fig. **5F**). These results indicate that COP1 directly interacts with and ubiquitinates RGL2 to induce its destabilization and suggest a molecular mechanism by which COP1 regulates RGL2 levels to induce seed germination.

### COP1 is as a positive regulator of germination-promoting genes that act downstream of *RGL2*

To inhibit germination, RGL2 represses the expression of *GASA6, EXPA* and *XTH* genes which promote seed germination (Zhong *et al*., 2015; Stamm *et al*., 2012; Rombolá-Caldentey *et al*., 2014; Yan *et al*., 2014). To further support a regulatory link between COP1 and RGL2, we investigated the effect of COP1 function on the expression of five germination-associated genes regulated by RGL2, including *GASA6*, *EXPA1*, *EXPA2*, *EXPA8*, and *XTH33* (Fig. **6**). In mock- and PAC-treatment conditions, these germination-associated genes were downregulated in *cop1-4* seeds and upregulated in *rgl2-SK54* seeds. In *cop1-4 rgl2-SK54* seeds, the expression levels of the germination-associated genes were quite similar to those in *rgl2-SK54* seeds, demonstrating that the negative effect of *cop1-4* mutation on their expression was cancelled out by the *rgl2* mutation. These results show that COP1, through RGL2 destabilization, positively regulates the expression of *GASA6* and *EXPA1*, as well as other genes related to cell wall remodeling that are involved in seed

**Figure 6.**
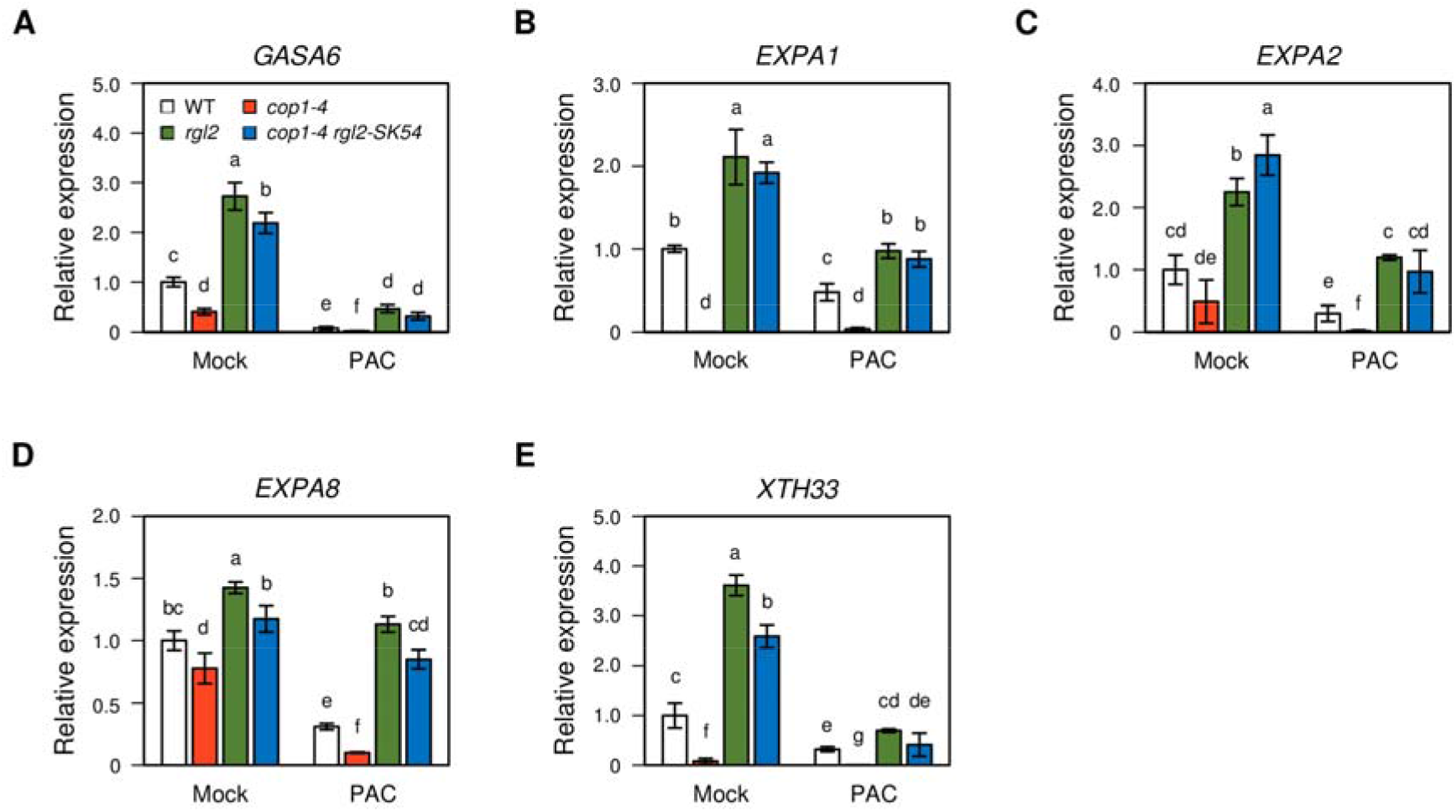
Effects of PAC on the expression of germination-associated genes in imbibed *cop1-4*, *rgl2-SK54*, and *cop1-4 rgl2-SK54* seeds. Altered expression of five *RGL2*-downregulated germination-associated genes, *GASA6*, *EXPA1, EXPA2, EXPA8,* and *XTH33*, in imbibed WT (Col-0), *cop1-4, rgl2*, and *cop1-4 rgl2-SK54* seeds in the presence or absence of 10 μM PAC at 3 DAI. The expression level of each gene obtained by qRT-PCR was normalized to that of *ACTIN2* (*ACT2*) and represented relatively to the expression levels in WT under normal conditions (mock), which is set as 1. The mean and SD were obtained from three biological samples (∼30 seeds/repeat). Different letters indicate significantly different values according to a one-way ANOVA and Duncan’s least significant range test (*P* < 0.05). DAI, day(s) after incubation to 22°C under LD; PAC, paclobutrazol.

### GA increases COP1 stability in the imbibed seeds

In imbibed seeds, increased GA biosynthesis is one of the most important triggers for the induction of seed germination (Steber *et al*., 1998). The increase in endogenous GA concentration decreases the stability of RGL2 (Tyler *et al*., 2004). Thus, we wondered whether GA or PAC have an impact on COP1 accumulation in imbibed seeds that could lead to changes in RGL2 stability. To examine the effects of GA and PAC on COP1 accumulation, we measured both *COP1* mRNA and COP1 protein levels in germinating seeds and during the onset of post-germinative seedling establishment after treatments with GA or PAC for 48 h (Fig. **7A-C**). The results showed that COP1 protein levels were increased by GA treatment 12 hours after imbibition and slightly decreased by PAC treatment (Fig. **7A-C**). The differences in COP1 protein accumulation in response to GA and PAC did not correlate with the variation pattern of *COP1* mRNA expression under these conditions, indicating that GA and PAC have post-translational effects on COP1 abundance (Fig. **7A-C**). These results suggest that increased GA levels upon seed imbibition promote a sustained accumulation of COP1 at the onset of the seedling establishment process. These results suggest that by regulating COP1 levels, GA promotes RGL2 degradation allowing seeds to germinate and contributing to attain adequate COP1 levels, required for seedling establishment and further development under light/dark cycles.

**Figure 7.**
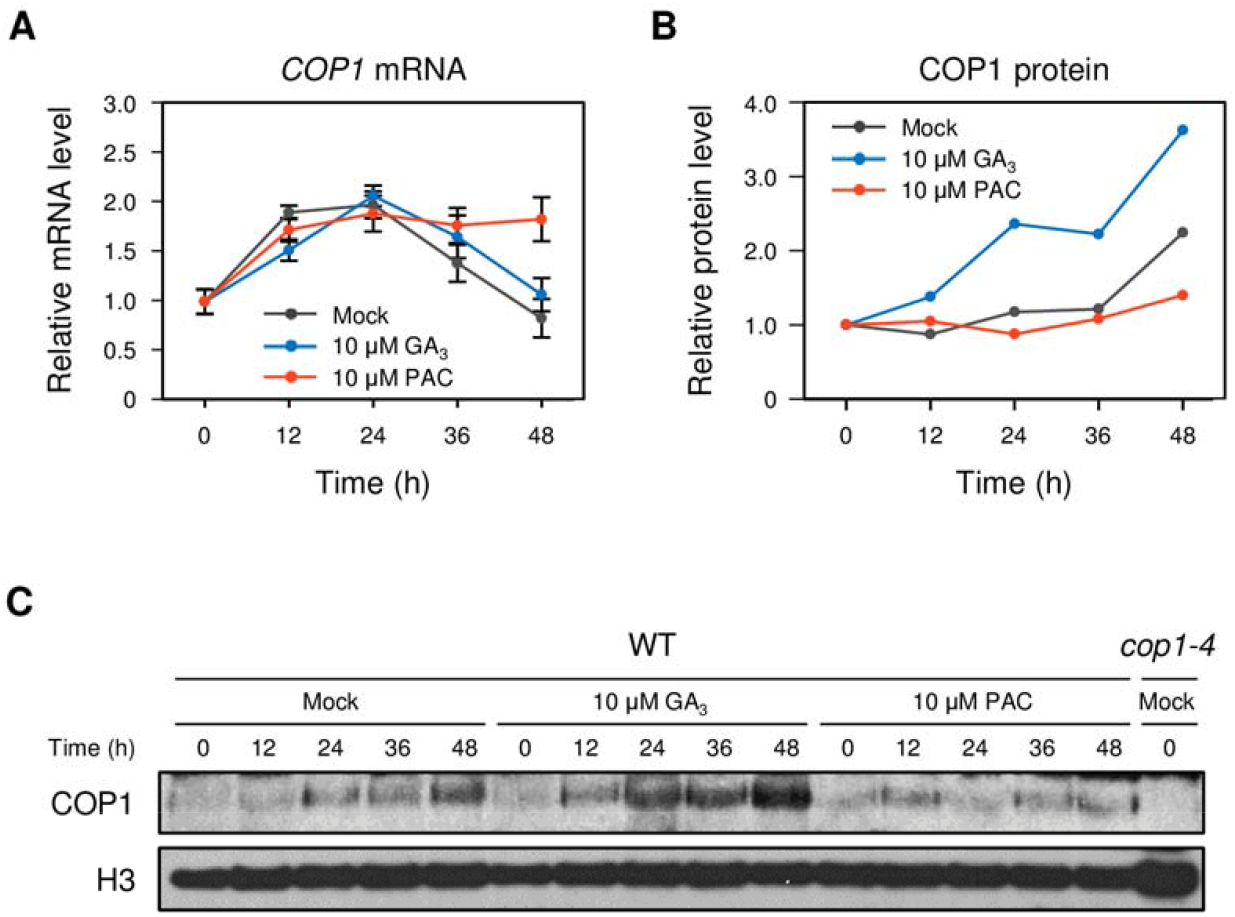
COP1 stability is enhanced by GA and decreased by PAC. (A) Relative expression levels of *COP1* in seeds stratified for 3 days at 4°C on MS medium and then transferred to 22°C. Seeds were sampled at the indicated time points after transfer. Relative expression levels of *COP1* were normalized to those of *ACTIN2* (*ACT2*). The expression levels of *COP1* are shown relative to the expression at time 0 under mock treatment, which is set as 1. In qRT-PCR analysis, the mean and SD were obtained from three biological repeats (∼30 seeds/repeat). (B, C) COP1 stability is increased by GA treatment but decreased by PAC treatment. COP1 and H3 histone protein levels were detected by immunoblot analysis and relative band intensity measured by ImageJ. Protein levels of each COP1 band were normalized to the level of H3 in each lane. The protein levels for each COP1 are shown relative to the expression at time 0 under mock treatment, which is set as 1. The mean and SD were obtained from three biological repeats (∼100 seeds/repeat). GA, gibberellin; PAC, paclobutrazol.

## Discussion

Seeds are equipped with molecular sensors to monitor surrounding environmental conditions and determine whether they are favorable for plant establishment (Seo *et al*., 2009). Here we describe a new regulatory module in which COP1 positively regulates seed germination by interfering with a component of the GA signaling pathway, the DELLA family member RGL2, which is a well-known repressor of seed germination. We showed that *cop1-4* mutants are strongly sensitive to PAC, while *COP1-OX/cop1-4* seeds are strongly insensitive to this GA biosynthesis inhibitor (Fig. **2**), mimicking the phenotype of the *rgl2-SK54* mutant (Fig. **4**, Fig. **S4**). In addition, COP1 promotes the degradation of RGL2 (Fig**. 5**). Supporting these findings, genetic analysis showed that *rgl2* is epistatic to *cop1*, as the double mutants show complete suppression of the *cop1-4* PAC hypersensitivity (Fig. **4A**) and of its defects on the activation of genes encoding cell wall proteins involved in cell loosening during germination (Fig. **6**). Furthermore, GA enhances COP1 protein accumulation upon imbibition (Fig. **7**), evidencing the GA’s broad role as a primary regulator in seed germination.

### COP1 regulates seed germination through RGL2

The CUL4^COP1-SPA^ E3 ubiquitin ligase is well described as a repressor of light signaling, targeting for degradation photomorphogenesis-promoting factors (Lau & Deng, 2012; Zhu *et al*., 2015). Our data show that *cop1* alleles display defects in seed germination that are unrelated to the fusca phenotypes as *cop10* fusca mutants do not display them. COP10 and DET1 belong to the same E3 ubiquitin ligase and it has been previously found that *det1* fusca mutant seeds germinate better than WT in normal conditions and are hypersensitive to ABA (Fernando and Schroeder, 2015). In the case of *cop1* mutant seeds, the defects can partially complemented by GA application and, likewise, can be exacerbated by the presence of PAC, suggesting that COP1 plays a role in seed germination by interacting with the GA signaling pathway (Fig. **1**). CUL4^COP1-SPA1^ complexes play a positive role in seed germination by promoting the rapid degradation of PIF1 under red and far-red light conditions (Zhu *et al*., 2015). Thus, by being degraded by COP1, PIF1 acts in light-mediated seed germination independently of GA, making unlikely that the COP1–PIF1 module could be responsible for GA-related germination defects. In addition, HY5, a well-described COP1 target in photomorphogenesis, is known to repress seed germination by activating *ABI5* gene expression in response to ABA and salt stress (Chen *et al*., 2008; Yu *et al*., 2016). Thus, HY5 role in seed germination seems to be unrelated with GA signaling. This is consistent with our observation that *pif1* and *hy5* seeds germinate similarly to wild-type seeds in the presence of PAC. Together with the fact that *pif1* has a null and *hy5* only a little suppressive effect on the *cop1* germination hypersensitivity to PAC, both transcription factors are unlikely to be involved in the GA-related seed germination response.

The best-described mutants showing PAC insensitivity during germination are *rgl2* mutants. Indeed, this characteristic is a specific signature of *rgl2* among other *della* mutants (Lee *et al*., 2002; Tyler *et al*., 2004; Cao *et al*., 2005) (Fig. **S4**). Moreover, among the five Arabidopsis DELLA proteins, RGL2 has been described as a primary player in seed germination. Upon GA perception by the GID1 receptor, GID-GA-DELLA complexes are degraded by SCF^SLY1^ in a GA-dependent manner, which is the canonical pathway for RGL2 degradation (McGinnis *et al*., 2003; Dill *et al*., 2004). According to our experiments, the *rgl2-SK54* mutation completely suppresses the germination defects of *cop1-4* seeds (Fig. **4**), suggesting that RGL2 acts downstream of COP1 in seed germination and might be a direct target of COP1 ubiquitin ligase activity, which would represent a novel mechanism for the targeted degradation of RGL2.

### COP1 destabilizes RGL2

We found that COP1 directly interacts with RGL2, in a mechanism involving SPA1, and mediates its ubiquitination to control RGL2 abundance (Fig. **5**). Thus, COP1-mediated degradation of RGL2 might occur in parallel with the canonical SLY1- and GA-dependent degradation. Blanco-Touriñan *et al*. (2020) recently reported that COP1-SPA1 complexes mediate the destabilization of two other DELLA proteins, GAI and RGA, to promote hypocotyl elongation in response to shade and temperature cues. These results are complementary to our findings, and suggest that the targeted degradation of DELLA proteins by COP1 goes far beyond germination and can be extended to other DELLA- and GA-regulated processes during plant development. A striking similarity between our results and those of Blanco-Touriñan et al. (2020) is that *in vivo*, COP1 requires the presence of SPA1 for interaction with GAI and RGA, though not for their ubiquitination (Blanco-Touriñan *et al*., 2020). Moreover, we report that an active COP1 protein with full capacity to dimerize and enter the nucleus is essential to maintaining wild-type germination levels in the presence of PAC (Fig. **S5**).

The CSN complex allows the activation of cullin-based E3 ubiquitin ligases by maintaining proper cycles of neddylation (an ubiquitin-like modification) and deneddylation of cullins. Previous studies have shown that CSN mutants have poor germination and hyperdormancy phenotypes (Wei & Deng, 2003; Dohmann *et al*., 2010; Franciosini *et al*., 2015; Jin *et al*., 2018). In the case of *csn1-10*, the hyperdormancy phenotype was totally dependent on the failure to degrade RGL2. This phenotypic defect might be due to altered neddylation states and activities of SCF^SLY1/2^ E3 ubiquitin ligases (Jin *et al*., 2018). However, it cannot be ruled out that *csn* mutations also impair the activity of CUL4^COP1-SPA^ complexes towards RGL2. Indeed, it has been recently shown that CRL4^CDDD^ complexes mediate COP1 degradation in a process that requires functional CSN (Cañibano *et al*., 2021). Supporting this notion, the deneddylation/neddylation ratios of CUL1, CUL3, and CUL4 have all been found to be higher after seed imbibition, suggestive of increased activity of CSN and various CULLIN-associated complexes during germination (Wei & Deng, 2003; Franciosini *et al*., 2015)

Our results uncovered a novel regulatory mechanism that restricts RGL2 function, suggesting that different mechanisms besides the canonical targeted degradation of RGL2 by SCF^SLY1/2^ act coordinately to govern seed germination. These mechanisms likely include GA-independent processes. In fact, the release from dormancy of *sly1* hyperdormant seeds is independent of the accumulation of the RGL2, GAI, and RGA DELLA proteins (McGinnis *et al*., 2003; Dill *et al*., 2004; Arizumi & Steber, 2007; Penfield *et al*., 2006). Indeed, *sly1* loss-of-dormancy germination correlates better with endogenous levels of ABI5 and seems to depend on ABA biosynthesis (Piskurevicz *et al*., 2008). These results highlight the complexity of the mechanisms involved in seed germination and release from dormancy.

### COP1 promotes the expression of cell wall modification genes and is induced by GA

RGL2 repression of seed germination depends on a number of transcription factors that end up connecting RGL2 function to structural genes that mechanically affect cell wall composition and the control of germination (Stamm *et al*., 2012; Rombolá-Caldentey *et al*., 2014; Yan *et al*., 2014; Sánchez-Montesino *et al*., 2019). For most target genes, their upstream regulatory mechanism is still unclear as is case for the *GASA6*–*EXP1A* regulatory module. GASA6 promotes cell elongation at the embryonic axis through the action of *EXPA1* by an unknown mechanism (Zhong *et al*., 2015). Since RGL2 represses *GASA6* and *EXPA1*. Thus, by repressing RGL2, COP1 can positively regulate the expression of these genes, supporting a role for COP1 in promoting embryonic axis cell elongation during seed germination (Figs. **6**, **8**). As shown by our analyses, COP1 promotes the expression of additional cell-wall-modifying genes that were previously reported to be targets of RGL2 in seed germination (Fig. **6**; Stamm *et al*., 2012).

**Figure 8.**
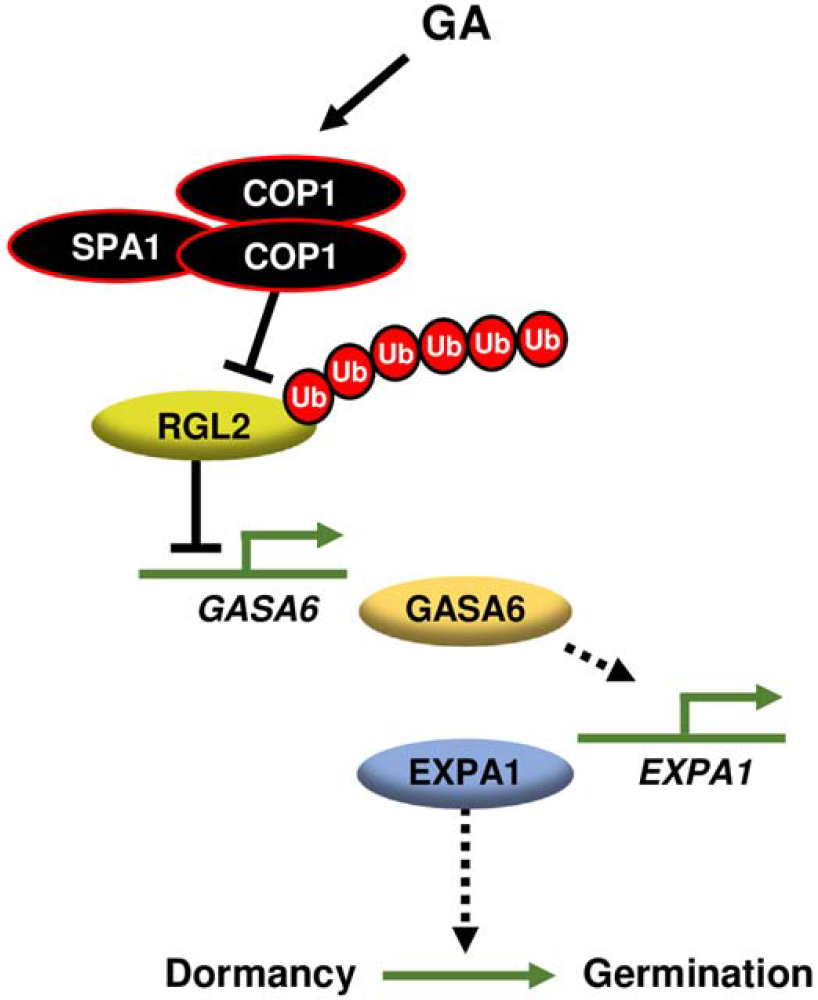
Model of the COP1–RGL2 regulatory module in seed germination. Upon perception of favorable environmental cues, GA is synthesized in the seeds to induce germination. *COP1* is expressed, and COP1 is stabilized by GA and interacts with RGL2, a negative regulator of the expression of germination-associated genes such as *GASA6*, *EXPA* genes, and *XTH*. The COP1–RGL2 interaction destabilizes RGL2, and consequently germination-associated genes are induced in the imbibed seeds. SPA1 is required for the *in vivo* interaction. Our model defines a non-canonical pathway by which GA inhibits RGL2 repression of seed germination through the activity of COP1. Arrows signify positive effects; blocked line, negative effect; dashed line, indirect regulation; GA, gibberellin; Ub, ubiquitin.

Notably, GA promotes COP1 protein accumulation during of seed germination and at the onset of seedling establishment (Fig. **7**). Though the mechanism behind this process requires further elucidation, it is clear that COP1 plays a major role in initial seedling development by promoting growth according to day/night cycles and circadian regulation contributing also for the ABA-mediated inhibition of post-germinative seedling establishment (Lau & Deng, 2012; Yadukrishnan *et al.,* 2020). In this way, increased accumulation of COP1 in response to GA might prevent precocious photomorphogenesis after seed germination and might afterward be necessary to maintain an equilibrium between the regulation of growth by elongation and photomorphogenic development in initial seedling developmental stages.

### Conclusions

Together, our data uncover a key role for COP1 in seed germination through promotion of the degradation of RGL2, a GA-regulated master repressor of seed germination. Therefore, COP1 contributes to the GA signaling pathway to promote seed germination and cell elongation, and thus is essential for initial seedling establishment. Further physiological and genetic studies will be key to fully understanding the GA-COP1 relations and fully integrating COP1 into the intricate network of seed germination regulatory components.

## Materials and Methods

### Plant materials and growth conditions

The *Arabidopsis thaliana* mutants used were of Columbia (Col) ecotype except for the *cop1-5* (Deng *et al*., 1992) and *cop10-1* (Wei *et al*., 1994) [Wassilewskija (Ws) ecotype] mutants, which are seedling lethal and were maintained as heterozygotes. The single mutants (Col ecotype) were *cop1-4* (a weak mutant allele; McNellis *et al*., 1994), *gai-t6* (Peng *et al*., 1997), *rga-28* (SALK_089146), *rgl1-SK62* (SALK_136162), *rgl2-SK54* (SALK_027654), *rgl3-3* (CS16355), *hy5-215* (Osterlund *et al*., 2000), and *pif1-1* (Oh *et al*., 2006). The double mutants used were *cop1-4 hy5-215* (Maier *et al*., 2013) and *pif1-1 cop1-4* (Xu *et al*., 2014). The *cop1-4 rgl2-SK54* double mutants were generated by crossing the single mutants and F2 genotyping with dCAPS (*cop1-4*; *Spe*I restriction enzyme digestion) and PCR (*rgl2-SK54*) primers (Table **S1**). For the *35S:COP1-GFP* (*COP1-OX*) constructs, the full-length *COP1* cDNA PCR amplified from Col-0 cDNA, cloned into the pDONR221 vector (Invitrogen) and subsequently into the pMDC85 plasmid (Curtis & Grossniklaus, 2003). Through *Agrobacterium tumefaciens* (GV3101)-mediated transformation in *cop1-4* or *rgl2-SK54* mutants by the floral-dip method (Clough & Bent, 1998) *COP1-OX/cop1-4* or *COP1-OX/rgl2-SK54* transgenic plants were obtained. The *COP1* mutant variants, i.e. the *35S:COP1*^WT^*-GFP/cop1-4, 35S:COP1*^L105A^*-GFP/cop1-4*, *35S:COP1*^L170A^*-GFP/cop1-4*, and *35S:COP1^cyt^-GFP/cop1-4* transgenic plants, were previously described (Lee *et al*., 2017).

### Germination rates

Fresh seeds (harvested within one month before the experiments) were used to measure germination rates. Seeds were surface sterilized with a solution containing 70% ethanol and 0.1% Triton X-100, for 20 min, and washed with 100% ethanol for three times. After being air-dried on sterile 3M filter paper, seeds were seeded on MS phytoagar medium (mock) or on the same medium supplemented with 10 μM GA_3_ or 10 μM PAC. For stratification, seeds were kept at 4°C in darkness for 72 h. Germination experiments were initiated with the transferred to a growth chamber at a constant 22°C temperature under cool white fluorescent light (100 μmol m^-2^ s^-1^) and long days (LD; 16-h light/day) and refered as days after incubation (DAI). To measure germination rates, germinated seeds were scored upon radicle emergence. For each experiment, three to five biological replicates of pools of about 100 seeds were used. Among each replicate seeds were collected from plants grown simultaneously under same conditions.

### Yeast two-hybrid (Y2H) assays

Y2H assays were performed using the Matchmaker GAL4 two-hybrid system (Clontech). The full-length and/or partial (RING; aa 1–104, CC; aa 121–213, WD-40 repeat; aa 371– 675) cDNAs of *COP1* were obtained by RT-PCR from wild-type (Col) plants (Yu *et al*., 2008) and cloned into the pGBK vector (as baits), and the full-length *RGL2* cDNA was cloned into the pGAD vector (as prey). Yeast (strain AH109) cotransformation was performed according to the Yeast Handbook (Clontech). An anti-HA (Roche) and an anti-myc (kindly provided by Xing Wang Deng) antibodies were used to check the expression of AD and BD fusion proteins.

### Bimolecular fluorescence complementation (BiFC) assays

The full-length cDNA of *RGL2* was gateway recombined to generate a YFC fusion into the BiFC plasmid sets (Belda-Palazón *et al.,* 2012). The YFN-COP1 and mRFP-SPA1 constructs were kindly provided by David Alabadi (Blanco-Touriñan *et al*., 2020). All the clones were transformed into *Agrobacterium tumefasciens* (GV301). Clones expressing fusion proteins as indicated were co-infiltrated into the abaxial leaf surface of 3-week-old *N. benthamiana* plants as described (Voinnet et al., 2003).The leaves were infiltrated with 50 μM MG132 the day previous to the observation. The p19 protein was used to suppress gene silencing. The empty vectors were used as negative controls. Fluorescence was visualized in epidermal cells of leaves after 3 d of infiltration using a TCS SP8 Leica Microsystems confocal laser microscope.

### Pull-down assays

For semi-*in vivo* pull-down assays, the full-length RGL2 coding sequence was cloned into the pKM596 (a gift from David Waugh, Addgene plasmid # 8837) and the MBP recombinant protein fusions were expressed in the *E. coli* BL21 (DE3). Recombinant proteins were purified and pull-down assays were performed according to Fonseca and Solano (2013). MBP-tagged protein fusions were purified using amylose agarose beads. Equal amounts of seedling protein extracts were combined with 10 µg MBP-tagged protein fusion or MBP protein alone, bound to amylose resin for 1 hr at 4°C with rotation, washed three times with 1 ml of extraction buffer, eluted and denatured in sample buffer before immunoblot analysis.

For *in vitro* pull-down assays, the full-length coding sequence of *RGL2* was cloned into the pGEX-4T-1 vector (Pharmacia) to generate a GST-RGL2 fusion protein, and transformed in the BL21-CodonPlus (Stratagene) *E. coli* strain. GST and GST-RGL2 were induced by IPTG, and purified using glutathione Sepharose resin beads (ELPIS Biotech, Korea) according to the manufacturer’s instruction. MBP and MBP-COP1 fusion protein were induced in BL21-CodonPlus (Stratagene) *E. coli* strain (Saijo *et al*., 2003) and purified using amylose resin beads (ELPIS Biotech, Korea). For in vitro pull-down assays, 2 μg of GST and GST-RGL2 proteins were incubated with immobilized MBP and MBP-COP1 proteins in binding buffer (50 mM Tris-HCl pH 8.0, 150 mM NaCl, and 1 mM EDTA) and incubated at 4°C for 2 h. After being washed three times with the binding buffer, the protein-retained beads were boiled in Laemmli buffer and immunoblotted using anti-GST and anti-MBP antibodies (Santa Cruz Biotechnology, USA).

### Immunoblotting on seed extracts

To detect endogenous COP1 protein levels, an anti-COP1 polyclonal antibody was used (Lee *et al*., 2017). Fresh seeds (harvested within 1 month before use) were imbibed in distilled water with or without 10 μM GA_3_ or 10 μM PAC, and harvested at each time point. Total crude extracts were prepared using extraction buffer (50 mM Tris-HCl pH 7.5, 4 M urea, 150 mM NaCl, 1 mM EDTA, and protease and phosphatase inhibitor mixtures (1 mM PMSF, 5 μg/mL leupeptin, 5 μg/mL aprotinin, 5 μg/mL pepstatin, 5 μg/mL antipain, 5 μg/mL chymostatin, 2 mM Na_2_VO_3_, 2 mM NaF and 50 μM MG132), separated by SDS-PAGE, and then immunoblotted with anti-COP1, anti-Myc (Santa Cruz Biotechnology, USA), and anti-GFP (Santa Cruz Biotechnology, USA) antibodies.

### Cell-free degradation assays

MBP-tagged RGL2 proteins were prepared from BL21-CodonPlus *E. coli* cells (Stratagene) and purified using an amylose resin according to the manufacturer’s instructions. For each reaction, 100 ng MBP-RGL2 or MBP proteins was incubated with 100 μg total soluble protein extracr at 22°C in assay buffer [50 mM Tris-HCl (pH7.5), 100 mM NaCl, 10 mM MgCl_2_, 5 mM DTT, and 5 mM ATP] from the wild-type (Col-0), *cop1-4*, and *COP1*-OX/*cop1-4* seedlings, previously grown at 22°C in the dark for 6 days. The reaction was stopped by adding Laemmli buffer at the respective times.

### Reverse transcription and quantitative real-time PCR (qRT-PCR)

Total RNA was extracted from seeds using the Fruit-mate (Takara, Japan) and MG RNAzol (Macrogen, South Korea) reagents according to the manufacturer’s instructions. First-strand cDNA was synthesized from 2 μg total RNA using M-MLV reverse transcriptase with oligo-dT primer (Promega). Expression levels of germination-associated genes were measured by qRT-PCR analysis using LightCycler 480 SYBR Green I Master mix (Roche) in a LightCycler 480 Real-Time PCR System (Roche, Basal, Switzerland). Expression levels were normalized by *ACTIN2* (*ACT2*). The gene-specific primer sets are shown in Table **S1**.

### *In vitro* ubiquitination assays

Assays were performed as previously reported (Yu *et al*., 2008) with minor modifications. Ubiquitination reaction mixtures contained 50 ng yeast E1 (Boston Biochem), 50 ng rice 6xHis-Rad6 (E2), 10 μg unlabeled ubiquitin (Boston Biochem), and 2 μg MBP-COP1 (previously incubated with 20 μM ZnCl_2_) in 30 μL of reaction buffer (50 mM Tris pH 7.5, 5 mM MgCl_2_, 2 mM ATP, and 0.5 mM DTT). As a substrate, 50 ng 6xHis-HA-RGL2-6His fusion was used per reaction. After 2 h incubation at 30°C, reactions were stopped by adding 30 μL of Laemmli buffer, and then half of each mixture (30 μL) was boiled for 5 min and separated by 7.5% SDS-PAGE. 6xHis-HA-RGL2-6His and its ubiquitinated conjugates were detected using anti-HA (1:1000; Roche) antibody.

### Accession numbers

*COP1*, At2g32950; *COP10*, At3g13550; *ACT2*, At3g18780; *GASA6*, At1g74670; *EXPA1*, At1g69530; *EXPA2*, At5g05290; *EXPA8*, At2g40610; *GAI*, At1g14920; *RGA*, At2g01570; *RGL1*, At1g66350; *RGL2*, At3g03450; *RGL3*, At5g17490; *XTH33*, At1g10550.

## Acknowledgements

We thank Xing-Wang Deng and Giltsu Choi for the *cop1-4, cop1-5, cop10-1, hy5-205, pif1-1, gai-t6, rga-28, rgl1-SK62, rgl2-SK54*, and *rgl3-3* mutant seeds and to Enamul Huq, Utte Hoecker and Giltsu Choi for the *pif1-1 cop1-4* and *cop1-4 hy5-215* mutant seeds, respectively. We thank to David Alabadi for providing the mRFP-SPA1 and YFN-COP1 clones used in BiFC assays. We are grateful to Yolanda Fernandez for technical assistance.

## Supplemental Data

**Supplementary Figure 1.** *35S:COP1-GFP* (*COP1-OX*) complements *cop1-4*.

**Supplementary Figure 2.** Epistasis analysis of *PIF1* and *COP1* in GA dependent seed germination.

**Supplementary Figure 3.** Epistasis analysis of *HY5* and *COP1* in GA dependent seed germination

**Supplementary Figure 4.** PAC-insensitive phenotype in *rgl2-SK54* mutant seeds.

**Supplementary Figure 5.** COP1 dimerization and nuclear localization are essential for PAC-insensitive germination in *COP1-OX/cop1-4* plants.

**Supplementary Figure 6.** Expression of *RGL2* and *COP1* in *cop1-4* and *rgl2-SK54* mutants, respectively.

**Supplementary Figure 7.** Expression of AD and BD protein fusions in Y2H experiments shown in Fig. 5A.

**Supplementary Table 1.** Primers used in this study.

## Notes

**Funding:** This research was supported by the Basic Science Research Program of the National Research Foundation (NRF) of Korea (NRF-2017R1A2B3003310 to N.-C.P.), Republic of Korea. S.F. has a Ramon y Cajal (RYC-2014-16308) grant funded by the Ministerio de Economia y Competitividad. Work in V.R.’s and and S.F.’s laboratory is funded by the Agencia Estatal de Investigación/Fondo Europeo de Desarollo Regional/European Union (BIO2016-80551-R; PID2019-105495GB-I00).

## Parsed Citations

Ariizumi T, Hauvermale AL, Nelson SK, Hanada A, Yamaguchi S, Steber CM. 2013. Lifting DELLArepression of Arabidopsis seed germination by nonproteolytic gibberellin signaling. Plant Physiology 162(4): 2125.

Ariizumi T, Steber CM. 2007. Seed germination of GA-insensitive sleepy1 mutants does not require RGL2 protein disappearance in Arabidopsis. Plant Cell 19(3): 791–804.

Belda-Palazón B, Ruiz L, Martí E, Tárraga S, Tiburcio AF, Culiáñez F, Farràs R, Carrasco P, Ferrando A. 2012. Aminopropyltransferases involved in polyamine biosynthesis localize preferentially in the nucleus of plant cells. PLoS One 7(10):e46907.

Bewley JD. 1997. Seed germination and dormancy. Plant Cell 9(7): 1055.

Blanco-Touriñán N, Legris M, Minguet EG, Costigliolo-Rojas C, Nohales MA, Iniesto E, García-León M, Pacín M, Heucken N, Blomeier T, et al. 2020. COP1 destabilizes DELLAproteins in Arabidopsis. PNAS 117 (24): 13792–13799.

Cañibano E, Bourbousse C, García-León M, Garnelo Gómez B, Wolff L, García-Baudino C, Lozano-Durán R, Barneche F, Rubio V, Fonseca S. 2021. DET1-mediated COP1 regulation avoids HY5 activity over second-site targets to tune plant photomorphogenesis. Molecular Plant. 14(6): 963–982. doi: 10.1016/j.molp.2021.03.009.

Cao D, Hussain A, Cheng H, Peng J. 2005. Loss of function of four DELLAgenes leads to light- and gibberellin-independent seed germination in Arabidopsis. Planta 223(1): 105–113.

Chen H, Zhang J, Neff MM, Hong S-W, Zhang H, Deng X-W, Xiong L. 2008. Integration of light and abscisic acid signaling during seed germination and early seedling development. Proceedings of the National Academy of Sciences 105(11): 4495–4500.

Clough SJ, Bent AF. 1998. Floral dip: a simplified method for Agrobacterium-mediated transformation of Arabidopsis thaliana. Plant Journal 16(6): 735–743.

Curtis MD, Grossniklaus U. 2003. Agateway cloning vector set for high-throughput functional analysis of genes in planta. Plant Physiology 133(2):462–9

Dill A, Sun T-P. 2001. Synergistic derepression of gibberellin signaling by removing RGAand GAI function in Arabidopsis thaliana. Genetics 159(2): 777–785.

Dill A, Thomas SG, Hu J, Steber CM, Sun TP. 2004. The Arabidopsis F-box protein SLEEPY1 targets gibberellin signaling repressors for gibberellin-induced degradation. Plant Cell 16(6): 1392–1405.

Dohmann EMN, Nill C, Schwechheimer C. 2010. DELLAproteins restrain germination and elongation growth in Arabidopsis thaliana COP9 signalosome mutants. European journal of cell biology 89(2-3): 163–168.

Earley KW, Haag JR, Pontes O, Opper K, Juehne T, Song K, Pikaard CS. 2006. Gateway-compatible vectors for plant functional genomics and proteomics. Plant J 45(4): 616–629.

Fernando VC, Schroeder DF. 2015. Genetic interactions between DET1 and intermediate genes in Arabidopsis ABA signalling. Plant Science 239:166–79.

Fonseca S, Solano R. 2013. Pull-down analysis of interactions among jasmonic acid core signaling proteins. In Jasmonate Signaling, pp. 159–171. Humana Press, Totowa, NJ.

Franciosini A, Moubayidin L, Du K, Matari Nahill H, Boccaccini A, Butera S, Vittorioso P, Sabatini S, Jenik Pablo D, Costantino P, et al. 2015. The COP9 SIGNALOSOME is required for postembryonic meristem maintenance in Arabidopsis thaliana. Molecular Plant 8(11): 1623–1634.

Fu X, Richards DE, Fleck B, Xie D, Burton N, Harberd NP. 2004. The Arabidopsis mutant sleepy1gar2-1 protein promotes plant growth by increasing the affinity of the SCFSLY1 E3 ubiquitin ligase for DELLAprotein substrates. Plant Cell 16(6): 1406–1418.

Gabriele S, Rizza A, Martone J, Circelli P, Costantino P, Vittorioso P. 2010. The Dof protein DAG1 mediates PIL5 activity on seed germination by negatively regulating GAbiosynthetic gene AtGA3ox1. Plant Journal 61(2): 312–323.

Holdsworth MJ, Bentsink L, Soppe WJ. 2008. Molecular networks regulating Arabidopsis seed maturation, after-ripening, dormancy and germination. New Phytologist 179(1): 33–54.

Jin D, Wu M, Li B, Bucker B, Keil P, Zhang S, Li J, Kang D, Liu J, Dong J, et al. 2018. The COP9 Signalosome regulates seed germination by facilitating protein degradation of RGL2 and ABI5. PLoS Genetics 14(2): e1007237.

Koornneef M, van der Veen JH. 1980. Induction and analysis of gibberellin sensitive mutants in Arabidopsis thaliana (L.) heynh. Theoretical and Applied Genetics 58(6): 257–263.

Lau OS, Deng XW. 2012. The photomorphogenic repressors COP1 and DET1: 20 years later. Trends in Plant Science 17(10): 584–593.

Lee B-D, Kim MR, Kang M-Y, Cha J-Y, Han S-H, Nawkar GM, Sakuraba Y, Lee SY, Imaizumi T, McClung CR, et al. 2017. The F-box protein FKF1 inhibits dimerization of COP1 in the control of photoperiodic flowering. Nature Communications 8(1): 2259.

Lee S, Cheng H, King KE, Wang W, He Y, Hussain A, Lo J, Harberd NP, Peng J. 2002. Gibberellin regulates Arabidopsis seed germination via RGL2, a GAI/RGA-like gene whose expression is up-regulated following imbibition. Genes and Development 16(5): 646–658.

Liu X, Hu P, Huang M, Tang Y, Li Y, Li L, Hou X. 2016. The NF-YC-RGL2 module integrates GAand ABA signalling to regulate seed germination in Arabidopsis. Nature Communications 7: 12768.

Maier A, Schrader A, Kokkelink L, Falke C, Welter B, Iniesto E, Rubio V, Uhrig JF, Hülskamp M, Hoecker U. 2013. Light and the E3 ubiquitin ligase COP1/SPAcontrol the protein stability of the MYB transcription factors PAP1 and PAP2 involved in anthocyanin accumulation in Arabidopsis. Plant Journal 74(4): 638–651.

McGinnis KM, Thomas SG, Soule JD, Strader LC, Zale JM, Sun TP, Steber CM. 2003. The Arabidopsis SLEEPY1 gene encodes a putative F-box subunit of an SCF E3 ubiquitin ligase. Plant Cell 15(5): 1120–1130.

McNellis TW, von Arnim AG, Deng XW. 1994. Overexpression of Arabidopsis COP1 results in partial suppression of light-mediated development: evidence for a light-inactivable repressor of photomorphogenesis. Plant Cell 6(10): 1391–1400.

Murase K, Hirano Y, Sun T-P, Hakoshima T. 2008. Gibberellin-induced DELLArecognition by the gibberellin receptor GID1. Nature 456(7221): 459.

Oh E, Kang H, Yamaguchi S, Park J, Lee D, Kamiya Y, Choi G. 2009. Genome-wide analysis of genes targeted by PHYTOCHROME INTERACTING FACTOR 3-LIKE5 during seed germination in Arabidopsis. Plant Cell 21(2): 403–419.

Oh E, Kim J, Park E, Kim J-I, Kang C, Choi G. 2004. PIL5, a phytochrome-interacting basic helix-loop-helix protein, is a key negative regulator of seed germination in Arabidopsis thaliana. Plant Cell 16(11): 3045–3058.

Oh E, Yamaguchi S, Hu J, Yusuke J, Jung B, Paik I, Lee H-S, Sun T-P, Kamiya Y, Choi G. 2007. PIL5, a phytochrome-interacting bHLH protein, regulates gibberellin responsiveness by binding directly to the GAI and RGApromoters in Arabidopsis seeds. Plant Cell 19(4): 1192–1208.

Oh E, Yamaguchi S, Kamiya Y, Bae G, Chung WI, Choi G. 2006. Light activates the degradation of PIL5 protein to promote seed germination through gibberellin in Arabidopsis. Plant Journal 47(1): 124–139.

Osterlund MT, Hardtke CS, Wei N, Deng XW. 2000. Targeted destabilization of HY5 during light-regulated development of Arabidopsis. Nature 405(6785): 462–466.

Park J, Lee N, Kim W, Lim S, Choi G. 2011. ABI3 and PIL5 collaboratively activate the expression of SOMNUS by directly binding to its promoter in imbibed Arabidopsis seeds. Plant Cell 23(4): 1404–1415.

Peng J, Carol P, Richards DE, King KE, Cowling RJ, Murphy GP, Harberd NP. 1997. The Arabidopsis GAI gene defines a signaling pathway that negatively regulates gibberellin responses. Genes and Development 11(23): 3194–3205.

Penfield S, Li Y, Gilday AD, Graham S, Graham IA. 2006. Arabidopsis ABA INSENSITIVE4 regulates lipid mobilization in the embryo and reveals repression of seed germination by the endosperm. Plant Cell 18: 1887–1899

Piskurewicz U, Jikumaru Y, Kinoshita N, Nambara E, Kamiya Y, Lopez-Molina L. 2008. The gibberellic acid signaling repressor RGL2 inhibits Arabidopsis seed germination by stimulating abscisic acid synthesis and ABI5 activity. Plant Cell 20(10): 2729–2745.

Ravindran P, Verma V, Stamm P, Kumar PP. 2017. ANovel RGL2-DOF6 Complex Contributes to Primary Seed Dormancy in Arabidopsis thaliana by Regulating a GATA Transcription Factor. Molecular Plant 10(10): 1307–1320.

Rombolá-Caldentey B, Rueda-Romero P, Iglesias-Fernández R, Carbonero P, Oñate-Sánchez L. 2014. Arabidopsis DELLAand two HD-ZIP transcription factors regulate GAsignaling in the epidermis through the L1 box cis-element. Plant Cell 26(7):2905–19.

Saijo Y, Sullivan JA, Wang H, Yang J, Shen Y, Rubio V, Ma L, Hoecker U, Deng XW. 2003. The COP1-SPA1 interaction defines a critical step in phytochrome A-mediated regulation of HY5 activity. Genes and Development 17:2642–2647.

Sanchez-Montesino R, Bouza-Morcillo L, Marquez J, Ghita M, Duran-Nebreda S, Gomez L, Holdsworth MJ, Bassel G, Onate-Sanchez L. 2019. Aregulatory module controlling GA-mediated endospermic expansion Is critical for seed germination in Arabidopsis. Molecular Plant 12(1): 71–85.

Seo M, Nambara E, Choi G, Yamaguchi S. 2009. Interaction of light and hormone signals in germinating seeds. Plant Molecular Biology 69(4): 463–472.

Shi H, Zhong S, Mo X, Liu N, Nezames CD, Deng XW. 2013. HFR1 sequesters PIF1 to govern the transcriptional network underlying light-initiated seed germination in Arabidopsis. Plant Cell 25(10): 3770–3784.

Shimada A, Ueguchi-Tanaka M, Nakatsu T, Nakajima M, Naoe Y, Ohmiya H, Kato H, Matsuoka M. 2008. Structural basis for gibberellin recognition by its receptor GID1. Nature 456(7221): 520–523.

Stacey MG, Hicks SN, Von Arnim AG. 1999. Discrete domains mediate the light-responsive nuclear and cytoplasmic localization of Arabidopsis COP1. Plant Cell 11(3): 349–363.

Stamm P, Ravindran P, Mohanty B, Tan EL, Yu H, Kumar PP. 2012. Insights into the molecular mechanism of RGL2-mediated inhibition of seed germination in Arabidopsis thaliana. BMC Plant Biology 12: 179–179.

Steber CM, Cooney SE, McCourt P. 1998. Isolation of the GA-response mutant sly1 as a suppressor of ABI1-1 in Arabidopsis thaliana. Genetics 149(2): 509–521.

Sun T-P. 2010. Gibberellin-GID1-DELLA: Apivotal regulatory module for plant growth and development. Plant Physiology 154(2): 567.

Tyler L, Thomas SG, Hu J, Dill A, Alonso JM, Ecker JR, Sun T-P. 2004. DELLAproteins and gibberellin-regulated seed germination and floral development in Arabidopsis. Plant Physiology 135(2): 1008.

Ueguchi-Tanaka M, Ashikari M, Nakajima M, Itoh H, Katoh E, Kobayashi M, Chow TY, Hsing YI, Kitano H, Yamaguchi I, et al. 2005. GIBBERELLIN INSENSITIVE DWARF1 encodes a soluble receptor for gibberellin. Nature 437(7059): 693–698.

Voinnet O, Rivas S, Mestre P, Baulcombe D. 2003. An enhanced transient expression system in plants based on suppression of gene silencing by the p19 protein of tomato bushy stunt virus. Plant Journal 33(5):949–56

Wei N, Chamovitz DA, Deng XW. 1994. Arabidopsis COP9 is a component of a novel signaling complex mediating light control of development. Cell 78:117–124.

Wei N, Deng XW. 2003. The COP9 signalosome. Annual Review of Cell and Developmental Biology 19:261–86.

Weitbrecht K, Müller K, Leubner-Metzger G. 2011. First off the mark: early seed germination. Journal of Experimental Botany 62(10): 3289–3309.

Wen CK, Chang C. 2002. Arabidopsis RGL1 encodes a negative regulator of gibberellin responses. Plant Cell 14(1):87–100.

Xu X, Paik I, Zhu L, Bu Q, Huang X, Deng XW, Huq E. 2014. PHYTOCHROME INTERACTING FACTOR1 enhances the E3 ligase activity of CONSTITUTIVE PHOTOMORPHOGENIC1 to synergistically repress photomorphogenesis in Arabidopsis. Plant Cell 26(5): 1992.

Yadukrishnan P, Rahul PV, Datta S. 2020. HY5 suppresses, rather than promotes, abscisic acid-mediated inhibition of postgermination seedling development. Plant Physiology 184(2): 574–578.

Yadukrishnan P, Rahul PV, Ravindran N, Bursch K, Johansson H, Datta, S. 2020. CONSTITUTIVELY PHOTOMORPHOGENIC1 promotes ABA-mediated inhibition of post-germination seedling establishment. Plant Journal, 103: 481–496.

Yan A, Wu M, Yan L, Hu R, Ali I, Gan Y. 2014. AtEXP2 is involved in seed germination and abiotic stress response in Arabidopsis. PLoS One 9: e85208.

Yu JW, Rubio V, Lee NY, Bai S, Lee SY, Kim SS, Liu L, Zhang Y, Irigoyen ML, Sullivan JA, et al. 2008. COP1 and ELF3 control circadian function and photoperiodic flowering by regulating GI stability. Molecular Cell 32(5): 617–630.

Yu Y, Wang J, Shi H, Gu J, Dong J, Deng XW, Huang R. 2016. Salt stress and ethylene antagonistically regulate nucleocytoplasmic partitioning of COP1 to control seed germination. Plant Physiology 170(4): 2340–2350.

Zhong C, Xu H, Ye S, Wang S, Li L, Zhang S, Wang X. 2015. Gibberellic acid-stimulated Arabidopsis6 serves as an integrator of gibberellin, abscisic acid, and glucose signaling during seed germination in Arabidopsis. Plant Physiology 169(3): 2288–2303.

Zhu L, Bu Q, Xu X, Paik I, Huang X, Hoecker U, Deng XW, Huq E. 2015. CUL4 forms an E3 ligase with COP1 and SPAto promote light-induced degradation of PIF1. Nature Communications 6: 7245.

